# Cost evaluation during decision making in patients at early stages of psychosis

**DOI:** 10.1101/225920

**Authors:** Anna O. Ermakova, Nimrod Gileadi, Franziska Knolle, Azucena Justicia, Rachel Anderson, Paul C. Fletcher, Michael Moutoussis, Graham K. Murray

## Abstract

Jumping to conclusions during probabilistic reasoning is a cognitive bias reliably observed in psychosis, and linked to delusion formation. Although the reasons for this cognitive bias are unknown, one suggestion is that psychosis patients may view sampling information as more costly. However, previous computational modelling has provided evidence that patients with chronic schizophrenia jump to conclusion because of noisy decision making. We developed a novel version of the classical beads-task, systematically manipulating the cost of information gathering in four blocks. For 31 individuals with early symptoms of psychosis and 31 healthy volunteers, we examined the numbers of ‘draws to decision’ when information sampling had no, a fixed, or an escalating cost. Computational modelling involved estimating a cost of information sampling parameter and a cognitive noise parameter. Overall patients sampled less information than controls. However, group differences in numbers of draws became less prominent at higher cost trials, where less information was sampled. The attenuation of group difference was not due to floor effects, as in the most costly block participants sampled more information than an ideal Bayesian agent. Computational modelling showed that, in the condition with no objective cost to information sampling, patients attributed higher costs to information sampling than controls (Mann-Whiney U=289, p=0.007), with marginal evidence of differences in noise parameter estimates (t=1.86 df=60, p=0.07). In patients, individual differences in severity of psychotic symptoms were statistically significantly associated with higher cost of information sampling (rho=0.6, p=0.001) but not with more cognitive noise (rho=0.27, p=0.14); in controls cognitive noise predicted aspects of schizotypy (preoccupation and distress associated with delusion-like ideation on the Peters Delusion Inventory). Using a psychological manipulation and computational modelling, we provide evidence that early psychosis patients jump to conclusions because of attributing higher costs to sampling information, not because of being primarily noisy decision makers.

## INTRODUCTION

A consistent psychological finding in schizophrenia research is that patients, especially those with delusions, gather less information before reaching a decision. This tendency to draw a conclusion on the basis of little evidence has been called a jumping to conclusions (JTC) bias (Garety & Freeman, 2013). According to cognitive theories of psychosis, a JTC bias is a trait representing liability to delusions. People who jump to conclusions easily accept implausible ideas, discounting alternative explanations, thus ensuring the persistence of delusions (Garety & Freeman 1999; Garety et al. 2007; Garety et al. 2013). Reviews and meta-analyses confirm the specificity, strength and reliability of the association of JTC bias and psychotic symptoms (Dudley, Taylor, Wickham, & Hutton, 2016; Fine, Gardner, Craigie, & Gold, 2007; P. A. Garety & Freeman, 2013; Ross, McKay, Coltheart, & Langdon, 2015; S. H. So, Garety, Peters, & Kapur, 2010; S. H. wai So, Siu, Wong, Chan, & Garety, 2015).

The JTC bias is commonly measured in psychosis research using variants of the “beads in the jar” task (Huq, Garety, & Hemsley, 1988). A person is presented with beads drawn one at time from one of two jars containing beads of two colours mixed in opposite ratios (P A Garety, Hemsley, & Wessely, 1991; Huq et al., 1988). Jumping to conclusions has been operationally defined in the ‘beads’ tasks as making decisions after just one or two beads (Garety et al., 2005; Warman, Lysaker, Martin, Davis, & Haudenschield, 2007). The commonly used outcome in this task is the number of beads seen before choosing the jar, known as draws to decision (DTD). The task requires the participant to decide of how much information to sample before making a final decision. This behaviour can be compared to the behaviour of an ideal Bayesian reasoning agent (Huq et al., 1988). However, when people experience evidence seeking as costly, it is thought that gathering less information could be seen as optimal leading to monetary gains at the expense of accuracy (Furl & Averbeck, 2011).

Although the JTC bias has been well replicated, the neurocognitive mechanisms underlying it, are unknown; many possible psychological explanations have been put forward, not all of which are mutually exclusive (Evans, Averbeck, & Furl, 2015). Motivational factors, such as intolerance of uncertainty (Bentall & Swarbrick, 2003; Broome et al., 2007), a “need for closure” (Colbert & Peters, 2002), a cost to self-esteem of seeming to need more information (Bentall & Swarbrick, 2003) or an abnormal “hypersalience of evidence” (Esslinger et al., 2013; Menon, Mizrahi, & Kapur, 2008; Speechley, Whitman, & Woodward, 2010) have been posited as potentially underlying the JTC bias. A common theme emerging from the intolerance of uncertainty, the need for closure, and the cost to self-esteem hypotheses is that patients experience an excessive cost of sampling information.

Computational models allow researchers to consider important latent factors influencing decisions. The “costed Bayesian model” (Moutoussis, Bentall, El-Deredy, & Dayan, 2011) incorporates a Bayesian consideration of future outcomes with the subjective benefits or penalty (cost) for gathering additional information on each trial and the noise during the decision making; Moutoussis and colleagues (2011) applied this computational model to the information sampling behaviour of a sample of chronic schizophrenia patients undertaking the beads task. They found, contrary to their expectations, that a higher perceived cost of the information gathering did not underpin the JTC bias. Therefore, they concluded that differences in the ‘noise’ of decision making were more useful in explaining the differences between patients and controls than the perceived cost of the information sampling. However, here we reason that a rejection of the ‘increased cost of information sampling’ account of the JTC bias based on this finding is premature, as patients with chronic schizophrenia may not be representative of all psychosis patients, especially not for those at early stages of psychosis - a stage particularly relevant for understanding the formation of delusions. A variety of different cognitive factors may contribute to the JTC bias; the balance of contributory factors may differ in different patient populations, with noisy decision making relating to executive cognitive impairments predominating in chronic, but perhaps not in early stages of psychosis. Furthermore, Moutoussis and colleagues (2011) applied their model to an existing dataset (Corcoran et al., 2008) which used the classic beads task. In this task, no explicit value was assigned to getting an answer correct or incorrect, and no explicit cost was assigned to gathering information. The authors themselves concluded that their work required replication, including incorporation of experimental manipulation of rewards and penalties.

The current study investigated the hypothesis that patients with early psychosis attribute higher costs to information sampling using a novel version of the traditional beads task and computational modelling. Focusing on patients at early stages of psychosis allowed us to investigate the JTC bias before the onset of a potential neuropsychological decline seen in some patients with chronic schizophrenia, and to study a largely unmedicated sample of psychosis patients. Specifically, we were interested in testing whether patients adapted their decision strategies when there is an explicit cost of information sampling. We, therefore, developed a variation of the beads task in which there were blocks with and without an explicit cost of information sampling, and we gave feedback for correct and incorrect answers. This manipulation allowed the comparison between groups on different cost schedules. However, it would also allow us to test the competing hypothesis, which is that psychosis patients jump to conclusions because of primarily noisy decision making behaviour. Under this account, patients should be insensitive to a cost manipulation in the novel setup of the paradigm and apply random decision making.

This study presents a novel investigation of the processes that lead to reduced information sampling in psychosis. We hypothesised that (1) psychosis patients would gather less information than controls when gathering information is cheap, and that (2) psychosis patients and control would adjust their information sampling according to experimental cost manipulations and that the adjustments would mitigate, but not abolish, the difference between the groups. Finally, we hypothesised that (3) the costed-Bayesian model applied to this paradigm and largely unmedicated early psychosis group would provide explanatory evidence for JTC bias in favour of less information sampling because of higher perceived costs rather than purely noisy decision making.

## METHODS

### Study participants

The study was approved by the Cambridgeshire 3 National Health Service research ethics committee. An early psychosis group (N=31) was recruited, consisting of individuals with first episode psychotic illness (N=14) or with At Risk Mental States (ARMS; N=17), from the Cambridge early intervention service in psychosis, CAMEO. Inclusion criteria were as follows: age 16-35 years, current psychotic symptoms meeting either ARMS or first episode of psychosis (FEP) criteria according to the Comprehensive Assessment of At-Risk Mental States (CAARMS) (Morrison et al., 2012; Yung et al., 2005). FEP patients met ICD-10 diagnostic Patients with FEP were required to meet ICD-10 criteria for a schizophrenia spectrum disorder (F20, F22, F23, F25, F28, F29) or affective psychosis (F30.2, F31.2, F32.3). Healthy volunteers (N=31) without a history of psychiatric illness or brain injury were recruited as control subjects. None of the participants had drug or alcohol dependence. Healthy volunteers have not reported any personal or family history of neurological or psychotic illnesses, and were matched with regard to age, gender, handedness, level of education, and maternal level of education. None of the patients with ARMS were taking antipsychotic medication, and four patients with FEP were on antipsychotic medication at the time of testing. All of the experiments were completed with the participant’s written informed consent.

### Behavioural “Jumping to Conclusions” Task

This was a novel task (Figure 1), based on previously published tasks (Garety et al., 1991; Huq et al., 1988) of reasoning bias in psychosis, but amended in the light of decision-making theory, according to which the amount of evidence sought is inversely proportional to the costs of information sampling. These costs include the high subjective cost of uncertainty, and the cost to self-esteem or other factors (Moutoussis et al., 2011). Participants were told that there are two lakes, each containing black and gold fish in two different ratios (60:40). The ratios were explicitly stated and displayed on the introductory slide. A series of fish was drawn from one of the lakes; all the previously ‘caught’ fish were visible, in order to reduce the working memory load. The participants were informed that fish were being ‘caught’ randomly from either of the two lakes and then allowed to ‘swim away’. We used a pseudorandomised order for each trial, which was the same for all participants. The lake where the fish were drawn from was also pseudorandomised.

**Figure 1:**
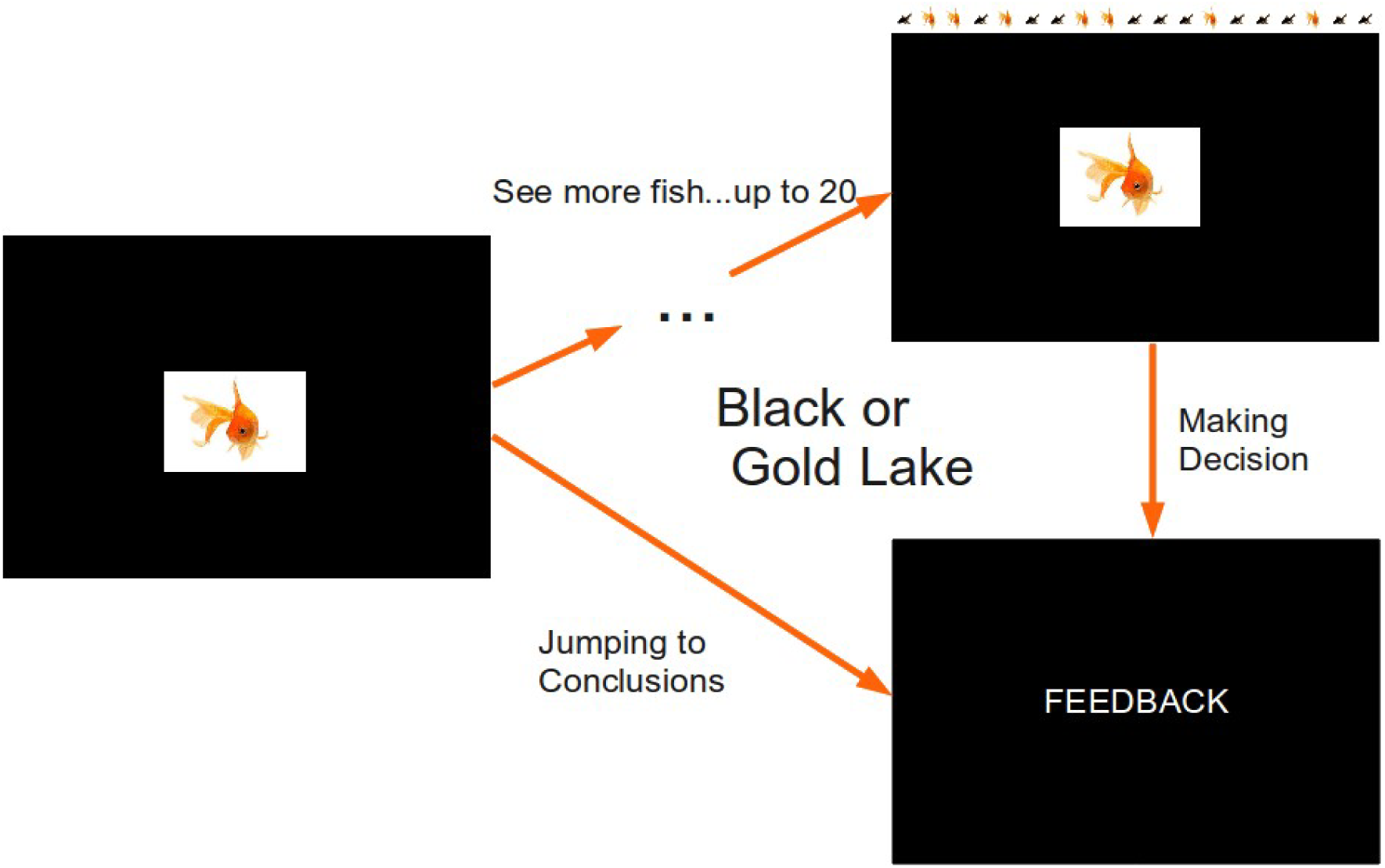
Experimental design of a single trial. In 50% of the trials fish were coming from the black lake, and in 50% they were coming from the gold lake. The order was pseudorandomised, so that the same sequences were used for all participants. Feedback, depending on the block, was either the words ‘Correct’ or ‘Incorrect’ in Block 1 or the number of points won or lost)during the trial in all subsequent blocks.

Participants could ask for a maximum of 20 fish to be shown. After each fish shown they either indicated whether the fish came from Lake G (gold) or B (black), or asked to see another fish. The trial terminated when the subject chose the lake. There were four blocks, each with the 10 trials of the predetermined sequences, in order to increase reliability. Block 1 was similar to the classical beads task, and was included to provide a reference point. The only difference was that feedback (‘correct’ or ‘incorrect’) was provided after each trial. In Block 2, a win was assigned to a correct decision (100 points) and a loss (−100 points) to an incorrect decision. In Block 3, the cost of each extra fish after the first one was introduced (−5 points) and was subtracted from the possible win or loss of 100 points for making a correct or incorrect decision respectively. Block 4 was similar to Block 3, but the information sampling cost was incrementally increased. The first fish would cost 0 points, the second −5 points, the third −10 points, etc.). Thus, the more fish would be sampled, the more points would be lost. Subjects performed the task at their own pace. Whether Lake G or B was correct was randomised. The task consisted of four blocks. Within each block there were 10 trials of predetermined sequences of fish, in order to increase reliability. All of the fish that were ‘caught’ during one trial were visible on the screen in order to minimise the working memory load. Block order was not randomised because the task increased in complexity.

The main outcome variable was the number of fish sampled (draws to decision, DTD). Secondary outcomes were the accuracy of the decision, calculated according to Bayes’ theorem based on the probability of the chosen lake given the colour and number of fish seen (Everitt & Skrondal, 2010), and the dichotomous JTC variable, which is defined as making decision after 2 or less pieces of information.

#### Partially observable Markov process decision-making

We consider a belief-based model of decision-making, formally a partially observable Markovian decision process (POMDP) to model behaviour in this task. The process is Markovian because we can concisely formulate the state that the people find themselves upon observing *n*_*d*_ draws, so that the state contains all information that can be extracted from observations thus far. As beads are drawn from one jar only at each trial, this can be simply defined as the number of *g* fishes seen so far, and the total number seen: *s* = [*n*_*d*_, *n*_*g*_]. The agent is interested to infer upon the true state of the world, which is a *B* or *G* lake. This is not directly observable, but ‘partially observable’. The agent maintains a belief component of their state, *P*(*G*|*s*). The Markov property still holds: future beliefs are independent of past beliefs given the current state (Fig. 2a). As we will see, belief-state transitions can be calculated just by considering the evidence so far. Given the current belief state, the probability of the possible unfolding of the task into the future can be estimated, and hence the expected returns for each possible future decision (Fig. 2b). The value of the available choices can thus be estimated: choosing the *B* lake, *D*_*B*_ the *G* lake *D*_*G*_, or sampling another piece of information, *D*_*s*_. The agent chooses accordingly, and either terminates the trial or gathers a new datum and repeats the process (Fig. 2).

**Figure 2.**
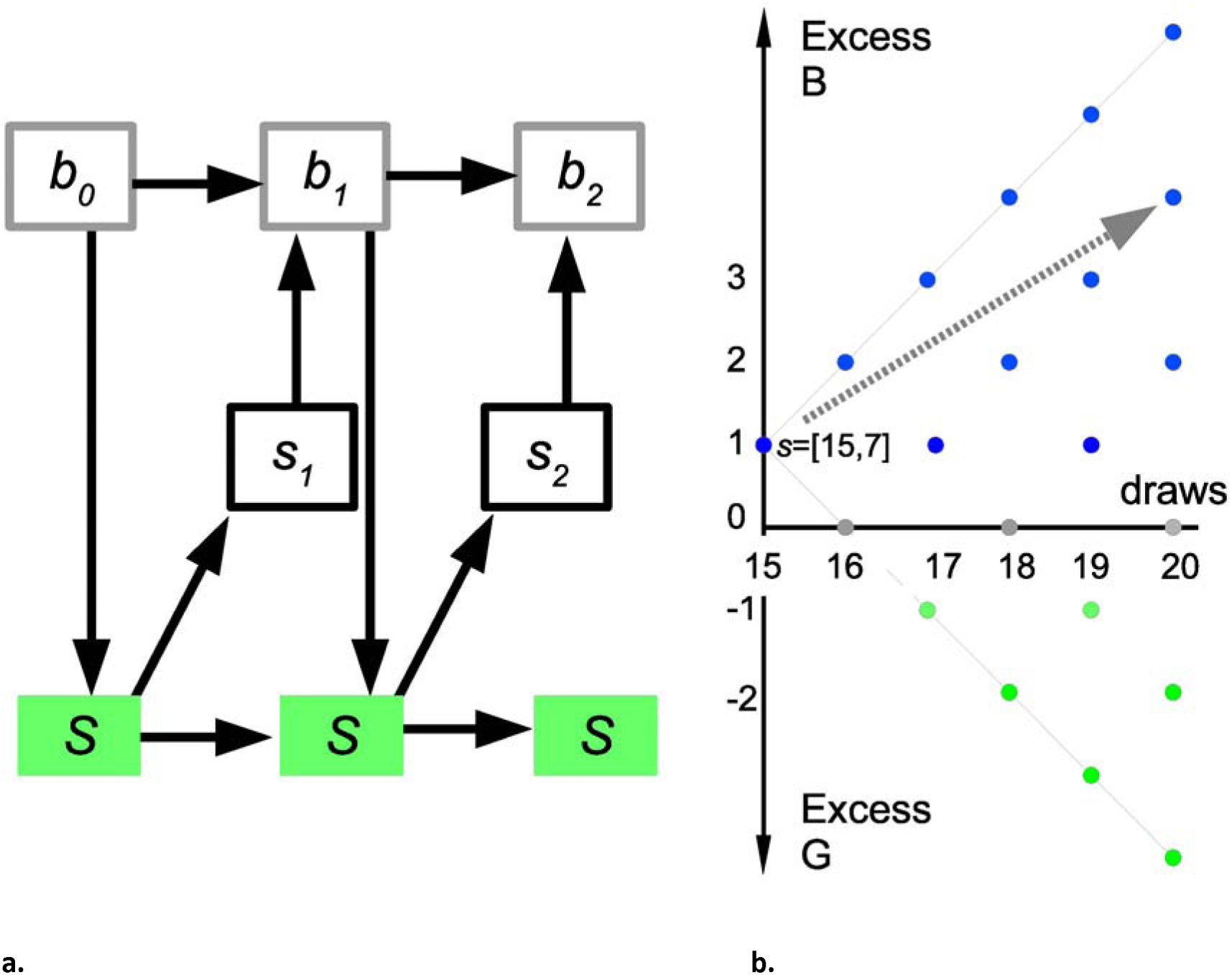
a. Markovian transitions in this task. Top: belief (probabilistic) component of states; middle: observable part of the state (data/feedback). Down arrows: actions (sample, declare). Bottom: true state. For example, let the cost of sampling be very high. Then *b*_*0*_ may be ‘equiprobable lakes’, action 1 ‘sample’, *s*_*1*_ ‘B’, *b*_*1*_ ‘60 % B’, action 2 ‘declare B’, and *s*_*1*_ ‘Wrong’. b. In this example, sampling cost is very low. A person has drawn 15 fishes, 7 of them *g.* The visible states corresponding to all possible future draws are shown. Looking ahead (example: grey arrow), the agent finds the ‘sampling’ action more valuable, in that the current preference for the *B* lake is likely to be strengthened at very low cost.

We now formally specify the model. Let *P*(*G* | *n*_*d*_,*n*_*g*_) be the probability of the lake being gold, after drawing *n*_*d*_ fish and seeing *n*_*g*_ gold fish. Using Bayes’ theorem, and assuming that a gold lake and a black lake are equally probable:

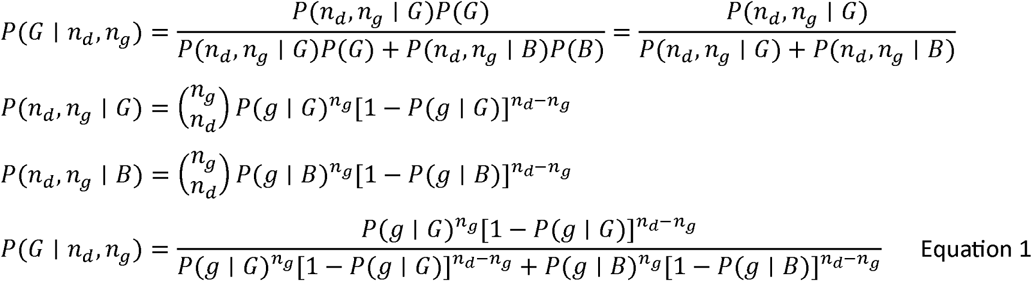

We then need to calculate the value of each action. For the ‘declare’ choices, the action value is the expected reward for correct answer, minus the expected cost for a wrong answer. For example, if *Q*(*D*_*g*_; *n*_*d*_,*n*_*g*_) is the action value of declaring the lake gold, after drawing *n*_*d*_ fish and seeing *n*_*g*_ gold fish,

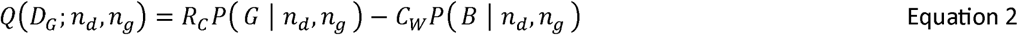

Where *R*_*C*_ is the reward for declaring the colour of the correct lake and *C*_*W*_ is the cost of declaring the colour of the wrong lake. In our task, *R*_*C*_ = *C*_*W*_, so:

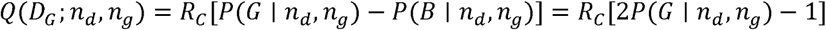

Similarly:

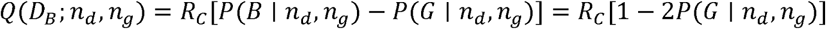

The action of sampling again is the expectation over the value of the next state, minus the cost of sampling *C*_*s*_(*n*_*d*_). If the value of the (possible) next state is *V*(*s*′), the action value for sampling again is the sum over the new possible states, weighted by their probabilities. The latter depends, in turn, on the identity of the true underlying lake, *L*:

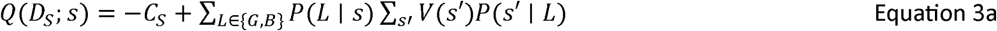

The possible outcomes for sampling again, and getting a black fish (going from (*n*_*d*_, *n*_*g*_) to (*n*_*d*_ + 1, *n*_*g*_)) and getting a gold fish (going from (*n*_*d*_, *n*_*g*_) to (*n*_*d*_ + 1, *n*_*g*_ + 1))

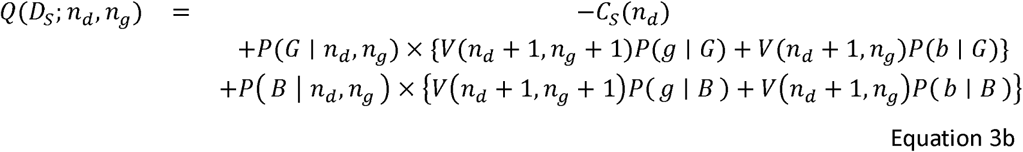

Agents will tend to prefer actions with the greatest value. An ideal, reward-maximizing agent, will always chose the action with the maximum value and will thus endow the corresponding state with this value. Denoting q as the vector of action values,

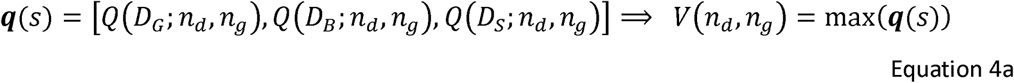

Real agents will choose probabilistically as a function of action-values, so:

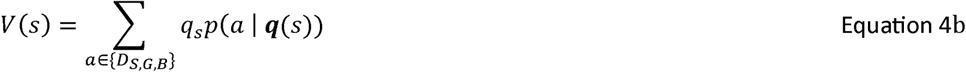

Agents cannot fill in the action values for sampling starting from their current state, as the next state value is not known. However, they can fill in all values by backward inference. At the very end, *n*_*d*_ = 20, sampling is not an option, and the action values can be calculated directly from Eq. 2 and the state value from Eq. 4. Once all possible state values for *n*_*d*_ *=* 20 have been calculated, Eqs. 3, 2 and 4 are used to calculate action and state values for *n*_*d*_ = 19, and downwards to *n*_*d*_ = 1.

We now turn to the ideal, deterministically maximizing agent against which we can compare human performance. When given the same sequence of fish as human participants, on average the ideal Bayesian agent samples 20, 20, 3.5 and 1 fish in each of the four blocks, respectively, achieving total winnings of 1070 points (Table 3). For no cost (Block 1 and 2), an ideal Bayesian agent samples all the fish and has p=0.835 of being correct (100% if we had infinite fish). For constant cost (5 points, Block 3), it samples until the difference between black and gold fish (Nb-Ng) is 2, with p=0.692 probability of being correct. For increasing cost (Block 4), it guesses after the first fish, so there is 0.6 probability of being correct. To model real agents, the probabilistic action choice in Eq. 4b took the softmax form:

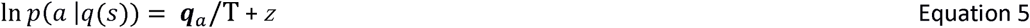

with *z* a normalizing constant same for all the actions at a specific state; and T a decision temperature parameter described below (Moutoussis et al., 2011).

Thus there are two parameters that shape a participant’s behaviour:

- CS: the cost of sampling, i.e. the subjective cost of each additional piece of information, compared to the final reward. It may be greater or smaller than the costs imposed by the experimenter, but is here taken to be constant (in a given block). High values of CS mean that decisions are made early.
- T: the noise parameter. As this value increases, the probability of the participant following the ideal behaviour (for their given value of CS) decreases, and their actions become more uniformly random.

#### Model parameter estimation

Here, we were interested in the most accurate possible estimates of the means and variances characterizing the psychosis and control groups. We, therefore, used what is known as a hierarchical model, a variant of the random effects approach. Here, a participant’s model parameters are drawn from a group, or population, distribution. This procedure estimates the mean and variance for CS and for T for the whole group. For a group of, for example, 30 participants with 10 data points each, we use 300 data points to estimate 4 values (i.e. mean and variance for each of the two parameter distributions), rather than 2 different parameters for each of the 30 participants (i.e. 60 parameters in total). We assumed both parameters to be positive, and a priori uncorrelated, and therefore appropriately modelled by independent gamma distributions. The standard ways that mathematicians parameterise gamma distributions are in terms of a ‘shape’ and ‘scale’ or ‘rate’. However, these do not map intuitively to the quantities we are usually interested in clinical research; that is, a measure of centre and a measure of spread. Thus, we follow Moutoussis et al. (2011) in describing our parameters by mean and variance. We used the technique expectation-maximization (EM, see Appendix; Dayan and Abbott, 2001; Moutoussis et al., 2011). In brief, EM proceeds by first assuming uninformative distributions at the group level; using these as uninformative priors, it derives probability distributions for the parameters of each participant based on each one’s data. This is ‘expectation’. Then, it re-estimates the group-level distributions to maximise the likelihood of the (temporarily fixed) lower level. This is ‘maximization’. The group-level distributions then form empirical priors that are used in place of the uninformative ones. The process is repeated until all estimates are stable; we ran between 25 and 30 iterations of the expectation maximisation algorithm (see Appendix). Runs took about 30s per iteration per participant on a single core of an Intel Core2 Duo CPU and 4GB of RAM.

Once the maximum likelihood parameters for a given group are estimated, they can be interpreted using the integrated Bayesian Information Criterion (iBIC (Huys et al., 2012, 2015)), which uses the likelihood values of best-fit parameters for two different models to decide which model best represents the data. In the case of the ordinary BIC, to assess model fit we calculate the maximum-likelihood of the data of each participant given a particular model, and we penalise this in proportion to the number of parameters in the model. The underlying assumption is that a greater number of parameters represents a proportionate reduction of prior belief that the participant belongs to a given region of the parameter space, and that all parameters are on the same footing for each participant. The volume of this parameter space scales to the power of its dimensionality, i.e. the number of parameters, so the log of this, used in the BIC, is proportional to this number of parameters. Redundant parameters, which would result in over-fitting, are thus penalised. However, this approximation can be refined. The study sample itself gives information about the prior probability that a particular parameter obtains at the micro-level, of the individual. This is the ‘empirical prior’, which, in our case, is calculated by expectation-maximization (Dayan and Abbot 2001). Now, the complexity penalty at the level of the individual is calculated by forming a mean, or integrated, likelihood weighed by this prior. This allows for the data to speak to some parameters being ‘more equal than others’ in penalizing complexity. We can now account well for the penalty due to the prior at the level of the individual, but we have not considered the level of the group. Should we assume that, say, patient participants should be fitted with different empirical priors than healthy controls, or the same? To compare separate-fit and common-fit statistical models, we turn to the BIC approximation of complexity proportional to the number of parameters, but now at the level of the groups. Thus, the ‘integrated BIC’, which is ‘integrated’ in the sense that it contains weighted-mean likelihoods rather than maximum-likelihoods, also contains a penalty term for models in proportion to the number of empirical priors, or groups, used. Therefore, it can be used for hypothesis testing: if one model has all participant parameters coming from one distribution (4 parameters), and a second model separates the control and unhealthy groups (8 parameters), a comparison of iBIC values can be used to decide if splitting the sample is justified. The interested reader is referred to Dayan and Abbot 2001, and Huys et al., 2012, 2015 for mathematical details.

In addition, we conducted analyses based on individual participant’s estimated model parameters. A reliable test of the hypothesis that the groups differ in the cost (or noise) parameters can be created by forcing the model to treat all participants as coming from one group, with a single group mean and variance, then using the model’s estimates of the single subject parameters to conduct a test of whether there are differences according to diagnostic group. This approach is over-conservative, but serves a purpose in subjecting the test of group differences to a stern challenge.

### Rating scales and questionnaires

The participants underwent a general psychiatric interview and assessment (Tables 1 and 2) using the Comprehensive Assessment of At Risk Mental States (CAARMS) (Morrison et al., 2012; Yung et al., 2005), the Positive and Negative Symptom Scale (PANSS) (Kay, Fiszbein, & Opler, 1987), the Scale for the Assessment of Negative Symptoms (SANS) (Andreasen, 1989) and the Global Assessment of Functioning (GAF) (Hall, 1995). The Beck Depression Inventory (BDI) (Beck, Steer, & Brown, 1996) was used to assess depressive symptoms during the last 2 weeks. IQ was estimated using the Culture Fair Intelligence Test (Cattell & Cattell, 1973). Schizotypy was measured with the 21-item Peters et al. Delusions Inventory (PDI-21) (Peters, Joseph, Day, & Garety, 2004).

**Table 1:**
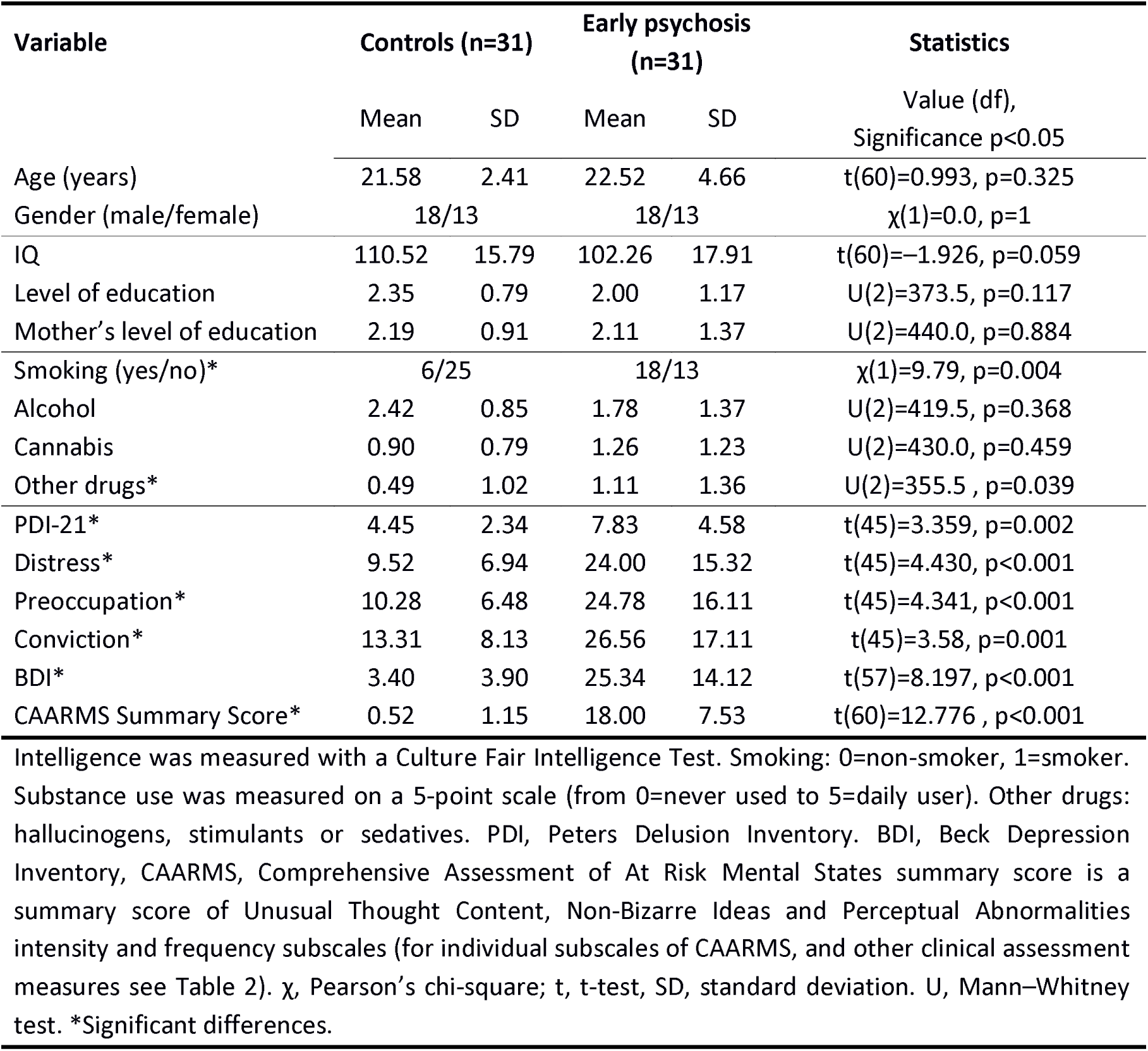
Sample characteristics for healthy controls and people with psychotic symptoms.

| Variable | Controls (n=31) |  | Early psychosis (n=31) |  | Statistics |
| --- | --- | --- | --- | --- | --- |
|  | Mean | SD | Mean | SD |  |
| Age (years) | 21.58 | 2.41 | 22.52 | 4.66 | t(60)=0.993, p=0.325 |
| Gender (male/female) | 18/13 | | 18/13 | | $\chi(1)=0.0$, p=1 |
| IQ | 110.52 | 15.79 | 102.26 | 17.91 | t(60)=-1.926, p=0.059 |
| Level of education | 2.35 | 0.79 | 2.00 | 1.17 | U(2)=373.5, p=0.117 |
| Mother's level of education | 2.19 | 0.91 | 2.11 | 1.37 | U(2)=440.0, p=0.884 |
| Smoking (yes/no)* | 6/25 | | 18/13 | | $\chi(1)=9.79$, p=0.004 |
| Alcohol | 2.42 | 0.85 | 1.78 | 1.37 | U(2)=419.5, p=0.368 |
| Cannabis | 0.90 | 0.79 | 1.26 | 1.23 | U(2)=430.0, p=0.459 |
| Other drugs* | 0.49 | 1.02 | 1.11 | 1.36 | U(2)=355.5, p=0.039 |
| PDI-21* | 4.45 | 2.34 | 7.83 | 4.58 | t(45)=3.359, p=0.002 |
| Distress* | 9.52 | 6.94 | 24.00 | 15.32 | t(45)=4.430, p<0.001 |
| Preoccupation* | 10.28 | 6.48 | 24.78 | 16.11 | t(45)=4.341, p<0.001 |
| Conviction* | 13.31 | 8.13 | 26.56 | 17.11 | t(45)=3.58, p=0.001 |
| BDI* | 3.40 | 3.90 | 25.34 | 14.12 | t(57)=8.197, p<0.001 |
| CAARMS Summary Score* | 0.52 | 1.15 | 18.00 | 7.53 | t(60)=12.776, p<0.001 |

**Table 2.** Clinical assessment measures for 31 patients with psychosis

|  | Mean | SD |
| --- | --- | --- |
| UTC | 2.44 | 2.19 |
| UTC frequency | 2.33 | 2.01 |
| NBI | 3.41 | 1.67 |
| NBI frequency | 3.52 | 1.48 |
| PA | 3.33 | 2.06 |
| PA frequency | 2.52 | 1.89 |
| DS | 0.59 | 1.28 |
| DS frequency | 0.96 | 1.99 |
| ADB | 1.81 | 1.82 |
| ADB frequency | 2.15 | 1.92 |
| SS | 1.59 | 1.48 |
| SS frequency | 1.30 | 1.44 |
| GAF score | 55.00 | 18.54 |
| SANS score | 0.35 | 0.75 |
| PANSS positive | 13.68 | 3.99 |
| PANSS negative | 9.87 | 4.88 |

BDI, Beck Depression Inventory; GAF, Global Assessment of Functioning. Comprehensive Assessment of At Risk Mental States (CAARMS) subscales: Unusual Thought Content (UTC), Non-Bizarre Ideas (NBI), Perceptual Abnormalities (PA), Disorganised Speech (DA), Aggression/Dangerous Behaviour (ADB), Suicidality and Self-Harm (SS). SANS, Scale for Assessment of Negative Symptoms

**Table 3.** Information sampling in controls, patients and an ideal Bayesian agent.

| Group | Control<br>n=31 | Psychosis<br>n=31 | Ideal<br>Bayesian<br>Agent |
| --- | --- | --- | --- |
|  | Mean (SD) | Mean (SD) |  |
| DTD 1 | 12.01<br>(5.42) | 8.14 (6.16) | 20 |
| DTD 2 | 12.69<br>(5.67) | 9.15 (6.31) | 20 |
| DTD 3 | 5.80 (4.23) | 4.14 (2.64) | Nb-Ng=+-2 |
| DTD 4 | 3.71 (3.12) | 3.03 (2.09) | 1 |
| P corr 1 | 0.72 (0.10) | 0.66 (0.10) | 0.835 |
| P corr 2 | 0.75 (0.09) | 0.69 (0.11) | 0.835 |
| P corr 3 | 0.65 (0.07) | 0.62 (0.08) | 0.692 |
| P corr 4 | 0.62 (0.09) | 0.59 (0.08) | 0.6 |

DTD, draws to decision (i.e. number of fish seen by the participant before reaching a decision); P corr, probability of being correct in Blocks 1 to 4; SD, standard deviation. Nb (number of black). Ng (number of gold).

### Statistical analysis of the behavioural data

The effect of manipulations of wins, losses and costs on the DTD and accuracy was assessed by repeated-measures analysis of variance (ANOVA) using SPSS 23 (IBM Corp.). Although DTD was not normally distributed, repeated-measures ANOVA is robust to violations of normality, and was therefore an appropriate test to run. IQ and depressive symptoms scores were initially included as covariates and dropped from the subsequent analysis where non-significant. We report two-tailed p-values, which were significant at p<0.05. When the assumption of sphericity was violated we applied Greenhouse–Geisser corrections. We also examined whether DTD was correlated with the severity of psychotic symptoms in the patient group (CAARMS positive symptoms) and with schizotypy scores on the PDI in controls using Spearman’s correlation coefficients. The intra-class correlation coefficients (ICCs) were used to estimate the consistency of decision making. We calculated the ICCs of the mean number of choices in each block separately for patients and controls.

For completeness and to help relate our study to prior literature (or for future meta-analyses), we report data on the dichotomous variable JTC, defined as making a decision after receiving one or two pieces of information, and we compare the FEP and ARMS patient groups separately to controls on Block 1 DTD and estimated model parameters.

## RESULTS

### Demographical characteristics of the participants

In Tables 1 and 2 the socio-demographic and cognitive parameters as well as clinical measures of the participants are presented. Although healthy volunteers had higher IQ, compared to the psychosis group, the difference was not significant, and the groups were matched with regard to level of maternal education and number of years of education. The control and patient group did not differ with regard to gender or age. There were significantly more smokers in the patient group compared to the control group, and some subjects of the patient group used more recreational drugs. For all of the other measures (e.g. alcohol or cannabis) there were no significant group differences. As expected there are significant differences between the healthy volunteers and the patients in all subscales of CAARMS (Table 2), on the self-reported depression questionnaire (BDI) and self-reported schizotypy scores (PDI). In the patient group only, we performed additional clinical assessments. The mean (±SD) score for PANSS positive symptoms was 13.68 (±3.99); for PANSS negative symptoms 9.87 (±4.88); SANS 0.35 (±0.75) and GAF 55.00 (±18.54).

### Group differences in the number of DTD and points

Inspection of Figure 3A reveals that in all of the blocks, the controls took more draws to decision than the patients. Mauchly’s test indicated that the assumption of sphericity was not violated (W(5)=0.211, p<0.001). On mixed-model ANOVA, there were significant main effects of block (F(3)=94.49, p<0.001 and of group (F(1)=5.99, p=0.017) on the number of DTD. The interaction between the group and the block was also significant (F(3)=4.32, p=0.006). This indicates that, depending on the group, block change had different effects on DTD. Group differences were statistically significant in the first two blocks (Block 1: p=0.007; Block 2: p=0.028), whereas in Block 3 and 4 there were the group differences were increasingly attenuated, as the cost of decision-making became increasingly high (Block 3: p=0.059; Block 4: p=0.419). Controls behaviour was more similar to the ideal Bayesian agent than patients on the first three blocks but not Block 4 (Table 3). Group differences in the probability of being correct were very similar to the results in the number of DTD (Figure 3B).

**Figure 3.**
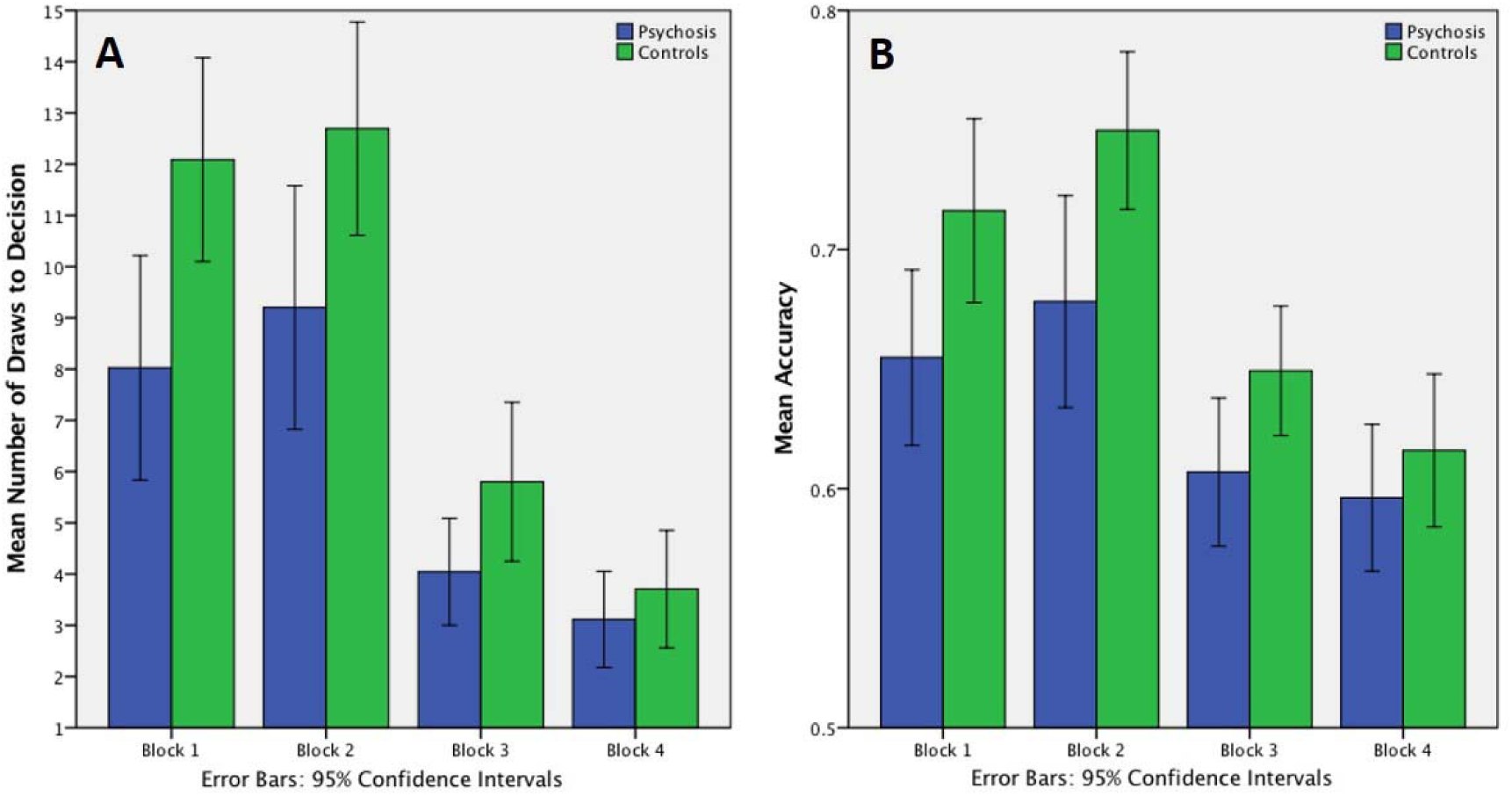
A) Mean number of draws to decision in the four blocks of the task. On mixed-model ANOVA, there were significant main effects of block (F(3)=94.49, p<0.001 and of group (F(1)=5.99, p=0.017). and an interaction between group and block (F(3)=4.32, p=0.006), with groups differences in block 1 (p=0.007) and block 2 (p=0.03). B) Probability of being correct (Accuracy) at the time of making the decision in four task blocks. Here there was an effect for block (F(2)=93.73, p<0.001), a marginally significant interaction between block × group (F(2)=2.52, p=0.086) and a significant groups effect (F(1)=4.14, p=0.047).

Analysing the points won/lost in Block 2-4 (Figure 4), we identified four outliers in each group that significantly exceeded ±2SD threshold. After excluding these subjects, the mixed-model ANOVA revealed a significant effect for block (F(2)=93.73, p<0.001), a marginally significant interaction between block × group (F(2)=2.52, p=0.086) and a significant groups effect (F(1)=4.14, p=0.047). Patients won significantly fewer points in Block 2 (F(2)=4.65, p=0.035), but did not differ from controls in Block 3 and 4 (both p>0.3).

**Figure 4:**
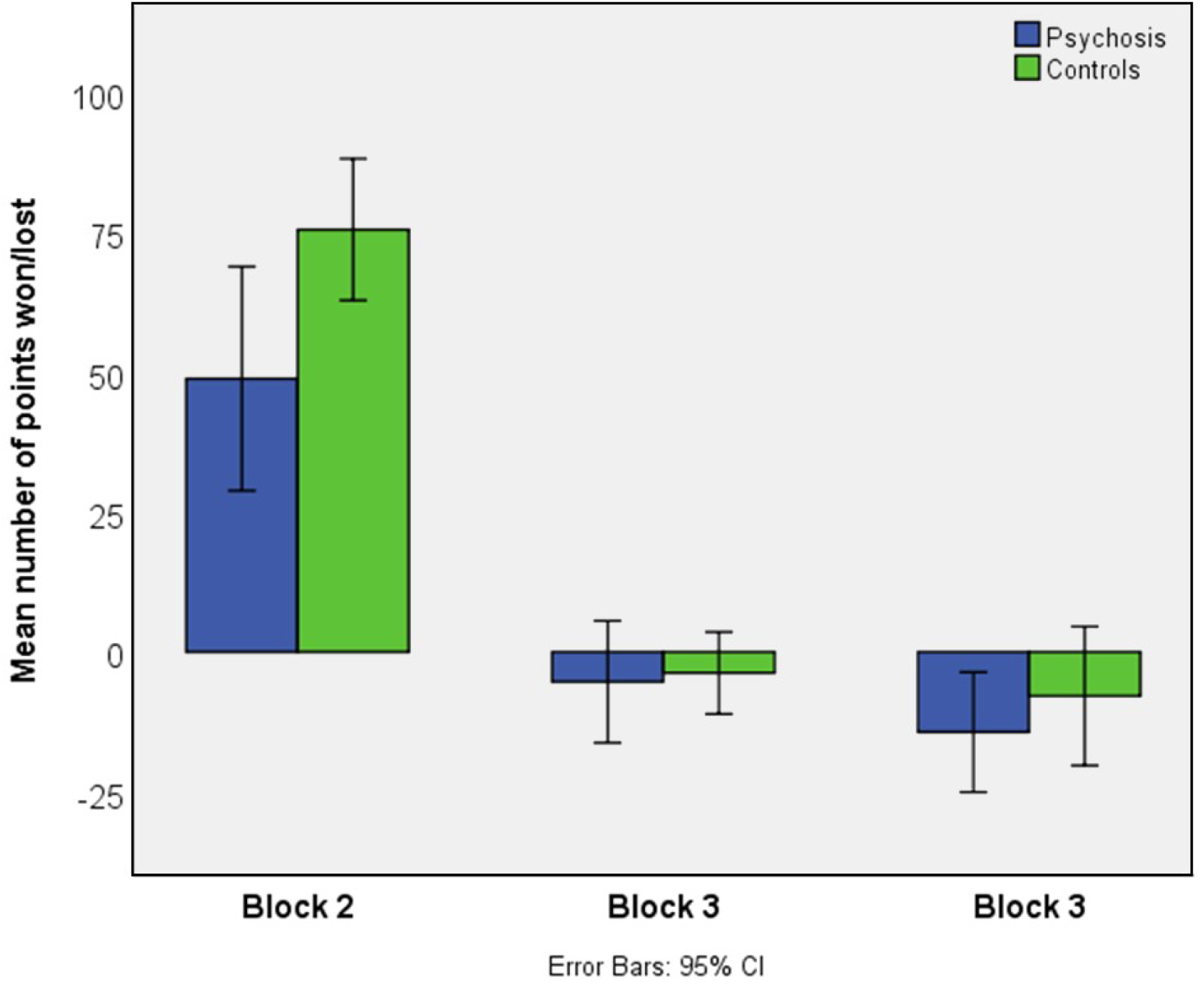
Mean number of point won or lost in Block 2-3 across all 10 trials. Patients won significantly fewer points in Block 2 (F(2)=4.65, p=0.035), but did not differ from controls in Block 3 and 4 (both p>0.3).

Percentage and count of people displaying a JTC reasoning style are presented in Table 4.

**Table 4.** Percentage and count of people who displayed JTC reasoning style.

| Group | Control n=31 |  | Psychosis n=31 |  |
| --- | --- | --- | --- | --- |
|  | Count | % | Count | % |
| JTC |  |  |  |  |
| No JTC | 18 | 58.1% | 15 | 48.4% |
| JTC Blocks 1/2 | 0 | 0.0% | 1 | 3.2% |
| JTC Blocks 3/4 | 11 | 35.5% | 7 | 22.6% |
| JTC all Blocks | 2 | 6.5% | 8 | 25.8% |

JTC (jumping to conclusions) is defined as reaching a decision after being given just one or two pieces of information.

### Intra-class correlations of DTD and SD of the mean DTD

Intra-class correlation coefficients (ICCs) of the mean number of DTD within each block were calculated separately for patients and controls. Within all blocks in both groups the correlations were very high, indicating that behaviour was consistent within each block (ICC values: Block 1: patients 0.965, controls 0.943; Block 2: patients 0.982, controls 0.973; Block 3: patients 0.972, controls 0.976; and Block 4: patients 0.973, controls 0.978).

### Correlation of symptoms and IQ with DTD

We calculated two-tailed Spearman’s correlations with positive psychotic symptoms in patients and with PDI scores in controls. In the patient group, we used a summary score of the three CAARMS subscales that quantify positive psychotic symptoms, namely Unusual Thought Content, Non-Bizarre Ideas (mainly persecutory ideas) and Perceptual Aberrations. An additional advantage of the summary measure is that it provides one measure to reduce the number of correlations that need to be performed. We also ran correlations BDI scores because the groups differed on this measure. In the group with psychotic symptoms the correlations with the overall CAARMS score were significant in the first three blocks (Block 1: rho=−0.515, p=0.003; Block 2: rho= −0.489, p=0.005; Block 3: rho= −0.491, p=0.005). There were no correlations in the psychotic symptoms group between DTD and BDI score (for all p>0.1).

In controls, we found significant correlations of DTD in Block 1 with the Distress (rho=−394, p=0.035) and Preoccupation (rho=−0.462, p=0.012) subscales of the PDI. DTD in Block 2 correlated with the Preoccupation subscale of PDI (rho=−0.387, p=0.038). There were no significant correlations with BDI scores (e.g. for Block 1 rho=−0.023, p=0.905).

In the patients, we furthermore found a positive correlation between IQ and the first three blocks (Block 1: rho=−0.481, p=0.006; Block 2: rho=−0.362, p=0.045; Block 3: rho=−0.364, p=0.044). There was no such correlation in the controls.

### Group differences in the probability of being correct (accuracy)

Mauchly’s test indicated that the assumption of sphericity has not been violated (W(5)=0.667, p<0.001). There was a significant main effect of block on the probability of being correct (F(3)=53.502, p<0.001) and a significant effect of group (F(1)=5.514, p=0.022). There was a significant interaction between the group and the block (F(1)=2.791, p=0.042), indicating that as the cost of decision making increased, the group differences became attenuated.

### Additional analyses having excluded participants on medication or to make groups more closely matched on IQ

In order to demonstrate that the findings were unaffected by antipsychotic medication, we conducted repeat analyses having excluded the four patients taking antipsychotic medication. The results were similar to in the full sample. As before, on mixed-model ANOVA, there were significant main effects of block (F(3)=88.59, p<0.001 and of group (F(1)=5.21, p=0.026) on the number of DTD. The interaction between the group and the block was also significant (F(3)=4.54, p=0.004). Group differences were statistically significant in the first two blocks (Block 1: p=0.013; Block 2: p=0.023), but reduced in Block 3 and 4 (Block 3: p=0.098; Block 4: p=0.55).

When we excluded the three highest IQ controls and two lowest IQ patients in order to make groups more similar in IQ (resulting in control mean IQ 107 and patient mean 105), the results were similar: on mixed-model ANOVA, there were significant main effects of block (F(3)=90.2, p<0.001 and of group (F(1)=5.25, p=0.026) on the number of DTD. The interaction between the group and the block was also significant (F(3)=3.50, p=0.017). Group differences were statistically significant in the first two blocks (Block 1: p=0.013; Block 2: p=0.039), but reduced in Block 3 and 4 (Block 3: p=0.062; Block 4: p=0.45).

### Computational Modelling Results: analysis of group estimates of model parameters

In our computational analysis of Block 1-3, we found that patients with early psychosis assigned a higher cost to sampling more data than healthy controls (Table 5). For example, in Block 1, where there was no explicit cost, the modelled mean cost of sampling in the controls was very low 1.9×10-3 compared to that of patients, 1.7. The modelled variance was also higher for the patient group (13) compared to the controls (2.0×10-6). The estimated noise parameters were similar in both groups. For example, in Blocks 1 and 2, the respective group estimated mean noise parameters were 3.4 and 3.6 in controls, and 4.2 and 2.9 in patients.

**Table 5:**
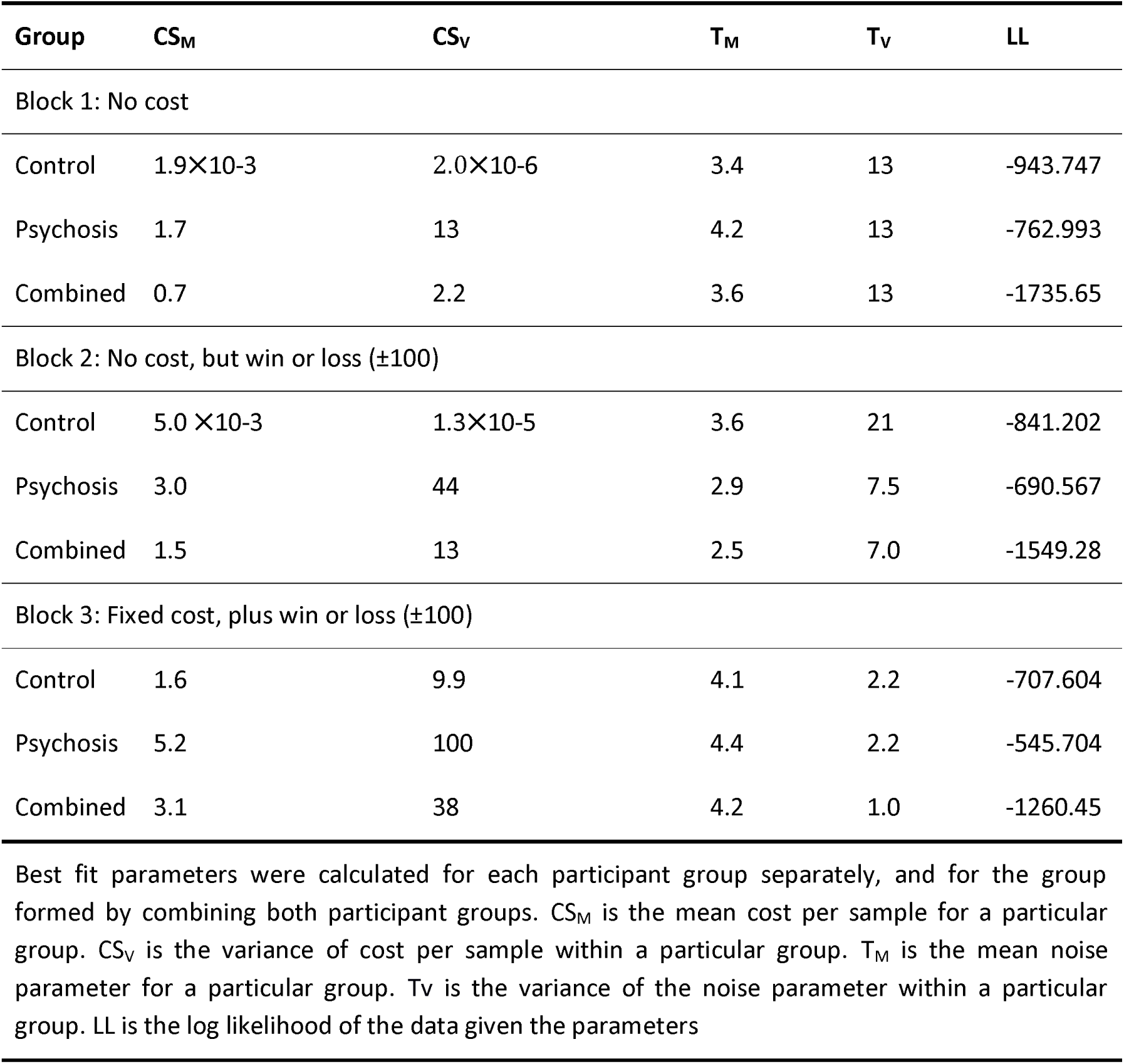
Best-fit distribution parameters for all groups in all experiments.

In Block 3, where an explicit cost of 5 points per fish sampled was assigned, the model shows that patients very slightly overestimated the cost of sampling compared to the assigned cost (estimated mean cost of sample for patients 5.2), whereas the controls underestimated it (estimated mean cost of sample 1.6).

In Block 4, we explicitly told participants that the cost for additional information is not fixed, but increases with amount of information requested. We did not fit the computational model to Block 4 because the model does not take into account the increasing cost structure set up.

To test the null hypothesis that both groups are drawn from the same distribution of behaviour parameters, we used iBIC. Table 6 shows the results of that calculation, where the null hypothesis can be rejected with strong evidence. For example, in Block 1, despite the fact the iBIC penalises the use of extra parameters substantially, when fitting CSm, CSv, Tm and Tv, separately for each group iBIC was 3450.2, compared to an iBIC of 3489.7 for fitting them as one combined group. Taken together, the iBIC can be used to compare the hypothesis about the two models of interest. In our case those are the following two: 1) unhealthy and healthy groups have independent CS and T parameter distributions, characterised by 8 numbers (mean and variance for CS and for T, for each group); and 2) all participants come from the same pool of noise and cost parameters, characterised by 4 numbers. The iBIC suggests that hypothesis 1 is better explained by the separate model in Blocks 1 and 2, and by the combined model in Block 3.

**Table 6:**
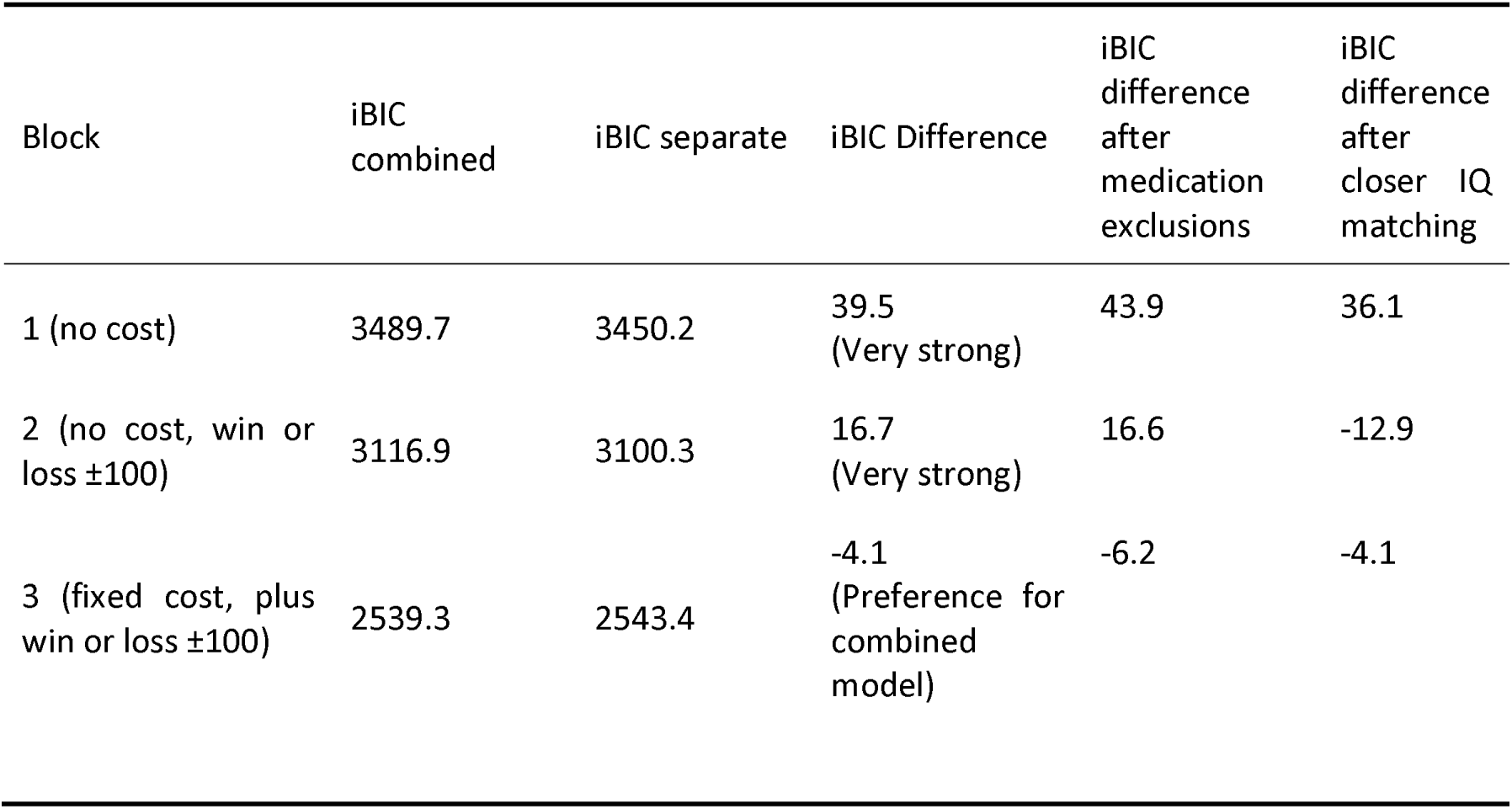
Integrated Bayesian Information Criterion (iBIC) values for the model where all participants are drawn from the same distribution, co pared to the model where the healthy and unhealthy groups differ in their distributions. iBIC values and differences are presented for analyses with all participants. iBIC differences are also shown for repeat analyses after exclusions for antipsychotic medication or closer IQ matching. Positive iBIC differences indicate the preference for separate groups, negative for a single combined group.

Model fits were not changed substantially after exclusion for medication (Table 6).

### Computational Modelling Results: analysis of individual participant level cost and noise parameters

A very strong test of the hypothesis that the groups differ in the cost (or noise) parameters can be created by forcing the model to treat all participants as coming from one group, with a single group mean and variance, then using the model’s estimates of the single subject parameters to conduct a test of whether there are differences according to diagnostic group. Although we found (see above) that in blocks 1 and 2 this assumption did not fit the data as well as modelling the groups separately, so the approach is over-conservative, this procedure serves a purpose in subjecting the test of group differences to a stern challenge. The groups differed significantly on estimated cost parameters in this procedure (block 1 controls mean=−0.24, median=−0.04, interquartile range=0.07, psychosis mean=−1.1, median=−0.1 interquartile range 1.2, Mann-Whiney U=289, p=0.007; block 2 controls mean=−0.8, median=−0.04, interquartile range=0.08; psychosis mean=−2.2, median=−0.14, interquartile range=4.9, U=363, p=0.038; block 3 controls mean=−2.0, median=−0.17, interquartile range=2.3, psychosis mean=−4.2; median=−0.48, interquartile range=8.62, U=338, p=0.045), but generally were similar on estimated noise parameters (block 1, controls mean=2.96, SD=2.08, patients mean 4.14 sd=2.86, t=1.86 df=60, p=0.07; block 2 controls mean=2.27, SD=1.61, patients mean =2.81, SD=1.95, t=1.2, p=0.23; block 3 controls mean= 4.1, SD=0.74, patients mean=4.31, SD=0.41, t=1.2 p=0.23).

The results were similar after exclusions for medication (Block 1 Cost group difference Mann-Whiney U=248, p=0.008; Noise group difference t=1.79 p=0.08), or to equalise IQ (Block 1 Cost group difference U=246, p=0.01; Noise group difference t=1.5, p=0.13).

Individually estimated greater cost parameters predicted higher psychotic symptom severity in patients (rho=0.58, p=0.001). There was no significant association between estimated noise parameters and psychotic symptom severity (rho=0.27, p=0.14). In controls, cost was associated with PDI preoccupation (rho = 0.41, p =0.03), and marginally with distress (rho 0.35, p =0.06), and noise was associated with overall PDI score (rho = 0.34, p =0.07), distress (rho = 0.42, p =0.02) and preoccupation (rho = 0.45, p = 0.01).

Cost parameter estimates were highly correlated across blocks: blocks 1 and 2 correlation coefficient rho=0.9; blocks 1 and 3 rho=0.6, and blocks 2 and 3 rho=0.7. Noise parameters were also correlated across blocks: noise on the first two blocks correlation co-efficient rho=0.8; noise on blocks 1 and 3 rho=0.3; noise on blocks 2 and 3 rho=0.4. However, in spite of associations across blocks, there were significant effects of block on both cost and noise. Repeated measures ANOVA effect of block on cost F=14, df=2,122; p=0.000003; effect of block on noise: F=22, df=2,122; p=5×10^-9^. Cost and noise parameters were related to each other (e.g. on block 1 rho=0.7), and, to a lesser extent, to IQ on some blocks (e.g. IQ versus block 1 cost rho=0.3, block 3 rho=0.3, block 1 noise rho= 0.2, block 3 noise rho=0).

### Subgroup analysis

On Block 1 DTD, controls (mean DTD 12.1 SD 5.4) gathered more evidence than FEP (mean 6.7, SD 6.0; one-tailed t=3.0, df=44, p=0.002), and ARMS (mean 9.2, SD=5.9; one-tailed t=1.7, df=47, p=0.045). On Block 1 cost parameter, controls (median 0.04, interquartile range 0.07) had lower values than FEP (median 2.5, interquartile range 4.1; Mann-Whitney U=106, one-tailed p=0.003) and ARMS (median=0.06, interquartile range 0.7, Mann-Whitney U=183, one-tailed p=0.04). On Block 1 noise parameter, controls (mean 3.0, SD=2.1) had lower values than ARMS (mean 4.2, SD 3.1; t=1.7, df=45, one-tailed T=0.045) and, marginally, than FEP (mean 4.0, SD 2.7; t=1.5, df=44, one tailed p=0.075)

## DISCUSSION

Our study shows that early psychosis patients generally gather less information before coming to a conclusion compared to healthy controls. Both groups slightly increased their DTD when rewarded for a correct answer (Block 2), and significantly decreased their DTD when there was an explicit cost for the sampling of information (Block 3 and 4). The decrease was strongest in Block 4 including the incremental cost increase for each ‘extra fish’ These effects were especially strong in controls, as they gathered significantly more information during Block 1 and 2 compared to patients. Thus, patients had a lower “baseline” against which to exhibit a change in DTD with increasing cost. However, we emphasise that an ideal Bayesian decision agent would still sample fewer information in than the average control or patient in Block 3 and 4 (in Block 4 the ideal Bayesian agent decides after the first fish, whereas our human participants sampled more). This indicates that potential floor effects may not be responsible for the decreased reduction of DTD in patients compared to controls. Together with our modelling results, this finding rather supports the hypothesis that independent of the objective cost-value of information, patients with early psychosis experience information sampling as more costly than controls.

In this novel version of the beads task using blocks with explicit costs, jumping to conclusions can be an advantageous strategy, because sampling a large amount of very costly information would cancel out the potential gain due to correctly identifying the lake. Consistent with this, group differences were especially strong in the first two blocks, where information sampling was free. In Block 3, when there was only a small cost of information, both groups responded to that change by lowering the number fish sampled. The difference between the groups was marginally significant on Block 3, where patients still applied fewer DTD (p=0.06). However, further increasing the information sampling cost completely abolished the group differences. Our data furthermore show that patients were significantly worse in overall accuracy (i.e. probability of being correct at the time of making the decision) and total points won indicating the application of an unsuccessful strategy. In general, these results show that healthy controls were more flexible in adapting their information sampling to the changed task blocks. Controls increased the number of fish sampled when there was a reward for the correct decision, and decreased it when information sampling had a cost. Patients also decreased the number of DTD when information sampling became more costly, but not as much as the controls, suggesting that the patients view information sampling generally as costly, somewhat independent of the actual value and the feedback. The results on Block 1 and 2, furthermore, indicate that psychosis patients have difficulties integrating feedback appropriately to update their future decisions. This is similar to results we reported in a recent study on the win-stay/lose-shift behaviour in a partially overlapping early psychosis group (Ermakova et al., 2018).

The variance in DTD across the two groups was similar and the ICCs were all greater than 0.94, indicating consistent decision making behaviour across the 10 trials in each block and within each group. If patients were acting more randomly, they would apply noisier and more variable decision making behaviour, but we did not observe this, which is in contrast to Moutoussis et al. (2011). Concluding from their Bayesian modelling data, they proposed that patients had more noise in their responses, leading to the reduced number of DTD. When modelling our data in a similar way, we found a difference between the groups in the perceived cost of information sampling, but less evidence in differences in the noise of decision behaviour. We suggest that the contrast between our findings and Moutoussis et al. (2011) may be due to the differences in the patient groups used in the two studies. Whereas Moutoussis et al. (2011) used chronic, mainly schizophrenia, patients with a potential neuropsychological decline and a lower IQ (of 92), our study used patients at early stages of psychosis with preserved cognitive functioning (IQ 102). Severity of psychotic symptoms was related to DTD and cost parameters in our sample; however, as all our participants were in the early stages of illness, we were not able to explore relationship with chronicity statistically. We did however, conduct secondary analyses to examine subgroup differences within our patient sample. On block 1 DTD and cost parameters, the patients with confirmed psychotic illness (first episode psychosis, FEP) had the most pronounced differences from controls, whereas regarding the noise parameter, there was little difference between the patient groups and indeed the ARMS group, with milder symptoms, had slightly higher noise parameters. Differences in task design may also contribute to differences in the results from Moutoussis et al (2011). Block 1 was most similar to that previous study: It differed in the type of test (computerised fishing emulation test versus the classic test using actual beads and actual jars), and in providing feedback (i.e. ‘correct’ or ‘incorrect’) after each decision, as well as in showing the sequence of fish drawn in each trial. These changes in our task potentially assisted patients (e.g. if some patients had memory deficits), which might also contribute to why we did not observe robust differences in the noise parameter in Block 1. When analysing the objective cost of information sampling in Block 3, we found that patients slightly overestimated the cost, while the controls underestimated it. Furthermore, key contributors to the best fit parameters may be those who jump to conclusions most: participants who consistently made a decision after viewing only one fish. The proportion of those individuals was significantly higher among the patients.

The percentage of people with psychosis who demonstrated JTC reasoning style in our study was relatively low, at 26%, compared to previous studies that report 40% of individuals (Dudley et al., 2011) or, half to two-thirds of the individuals with delusions (S. H. So et al., 2010). Likewise, our objective DTD values were slightly higher than that in most other studies. This might be due to the use of early stage psychosis patients and presence of feedback in our task compared to previous studies. Feedback has been shown to increase information sampling and accuracy both in patients with delusions and in controls (Lincoln, Ziegler, Mehl, & Rief, 2010).

Additionally, we found a correlation between IQ and DTD in the first three blocks in the patient group, so we cannot completely exclude the contribution of intelligence to the information-gathering bias. Some argue that impaired executive functions or working memory deficits contribute to the JTC bias (Falcone et al., 2015; P. Garety et al., 2013). A low IQ could lead to a low tolerance for uncertainty and an equivalent high cost of the information sampling, as well as the inability to integrate feedback in order to update future decisions. If more information is unpleasant, because it exceeds one’s capacity to utilise it, it could be viewed as costly. However, the fact that the correlation between IQ and DTD does not appear in controls argues against this. The groups were well matched on maternal education level (a proxy for premorbid or potential IQ) but the patient group had lower current IQ than controls (as expected given the that schizophrenia spectrum disorders are robustly associated with reduced current IQ compared to the general population). When we excluded the three controls with the highest IQ and the two patients with the lowest IQ, the DTD results were broadly unchanged. Computational modelling in block 1 was not changed by these exclusions, although in block 2 there were some differences. After the exclusions, in block 2, the BIC values did not suggest evidence that the participants from different diagnostic groups are drawn from different populations. However, when we examined the individual level modelled parameters, even after the exclusions there was evidence that patients had higher estimated sampling costs compared to controls. Taken together, the findings indicate that lower IQ associated with psychosis is likely to contribute to the JTC bias, but is unlikely to solely explain its existence in psychosis. The results of the study were unchanged when we excluded four patients taking antipsychotic (dopamine receptor antagonist) medication, which is consistent with two prior studies in healthy volunteers suggesting that dopaminergic manipulations do not have a large effect on information sampling (Andreou et al 2013, Ermakova et al 2014).

When comparing our results to those of Moutoussis et al (2011), the use of the same computational model, implemented in the same way, is advantageous. However, we note that there are limitations in the approach. For example, the use of a gamma distribution may not be optimal in the case of values near zero (Moutoussis et al 2011). It could be hypothesised that the degree of cognitive noise should be a constant per individual, and that thus it would be more parsimonious to apply the same noise parameters for a given subject across blocks. Our data suggest this is not the case. Noise is highly correlated across blocks, just as cost is. However, the experimental manipulation of block had a highly significant effect on both cost (as intended by our paradigm design) and noise (an incidental effect). Noise parameters were reduced in block 2, where a correct decision is explicitly rewarded, compared to block 1, where it is not (indicating that information sampling is not immutable but can be adaptively altered by psychological manipulation). Noise parameters were greater in block 3 than block 2, presumably because the decision is more difficult in block 3 (where participants need to balance the stated benefits and costs of sampling). When decisions reach a certain level of difficulty, participants may appear more random in their decision making because they can no longer effectively utilise the information available.

Rather than focusing on ICD-10 or DSM-V schizophrenia patients, we studied a group of patients early in their course of psychosis. All patients in our study suffered from current psychotic symptoms. Psychosis is often viewed as an upper part of the continuum, ranging from rare occurrences of delusions or hallucinations at one end, through individuals with regular ‘schizotypal’ traits (van Os, Linscott, Myin-Germeys, Delespaul, & Krabbendam, 2009). Consistent with this approach, and with the theory that a jumping to conclusions style cognitive bias contributes to psychotic symptom formation (Huq et al 1988), we found that patients with more severe positive symptoms sampled less information and had higher estimated sampling cost parameters. To investigate the idea of the psychosis continuum further, we looked at the correlations between information sampling and schizotypy characteristics in healthy volunteers. In this group, we found a negative correlation between the number of DTD and the scores on the distress and preoccupation subscales of the PDI, indicating that less information sampling is associated with higher scores. This is consistent with studies by Colbert and Peters (2002) and Lee et al. (2011), as well as the recent meta-analysis by Ross et al. (2015), in keeping with a continuum model of psychosis. Regarding modelled parameters in controls, estimated noise and was associated with total PDI score, PDI distress and PDI preoccupation, and estimated sampling costs were associated with PDI preoccupation, and marginally, with distress. This hints at the intriguing possibility that hasty decision making due cognitive noise may be a more important contributory factor to delusion-like thinking in the healthy population than it is to psychotic symptoms in psychotic illness, where hasty decision making due to higher information sampling costs appear to be more important.

### Summary

In summary, we found that early psychosis patients demonstrate a hasty decision-making style compared with healthy volunteers, sampling significantly less information. This decision-making style was correlated with delusion severity, consistent with the possibility that it may be a cognitive mechanism contributing to delusion formation. Our data are not consistent with the account that patients sample less information because they are in general more noisy decision makers. Rather, our data suggest that patients with psychosis sample less information before making a decision because they attribute a higher cost to information sampling. Although psychosis patients were less able to adapt to the changing demands of the task, they did alter their decision making style in response to the changing explicit costs of information, indicating that an impulsive decision making style is not completely fixed in psychosis. This finding is consistent with the possibility that information sampling may be a treatment target, e.g. for psychotherapy (Moritz et al., 2014), and that patients with psychosis may benefit in this neuropsychological domain, as they have in other domains, from cognitive scaffolding approaches exemplified in cognitive remediation therapy (Cella and Wykes 2017).

## Acknowledgments, funding and disclosures

Supported by a MRC Clinician Scientist [G0701911] and an Isaac Newton Trust award to GKM; by the University of Cambridge Behavioural and Clinical Neuroscience Institute, funded by a joint award from the Medical Research Council [G1000183] and Wellcome Trust [093875/Z/10/Z]; by awards from the Wellcome Trust [095692] and the Bernard Wolfe Health Neuroscience Fund to PCF; by the Cambridgeshire and Peterborough NHS Foundation Trust and Cambridge NIHR Biomedical Research Centre, and by The Max Planck - UCL Centre for Computational Psychiatry and Ageing, a joint initiative of the Max Planck Society and University College London. MM also receives support from the UCLH Biomedical Research Centre and is funded staff in the ‘Neuroscience in Psychiatry Network’, Wellcome Strategic Award (095844/7/11/Z).

PCF has consulted for GlaxoSmithKline and Lundbeck and received compensation. NG is employed by Google but this work was not part of his employment there and he did this work in his spare time. FK, AOE, GKM, MM, and AJ have no conflicts of interest.

## Author roles

Author roles: AE: Conceptualization, methodology, formal analysis, writing (original draft preparation, review and editing); NG. methodology, formal analysis, writing (review and editing); FK (methodology, formal analysis, writing (review and editing); AD. Methodology, investigation, project administration, writing (review and editing); RA (methodology, formal analysis, writing, review and editing); PCF Conceptualization, methodology, supervision writing (review and editing); MM methodology, analysis supervision, writing (original draft preparation, review and editing); G.K.M., conceptualization, project administration, methodology, formal analysis, supervision, funding acquisition, writing (original draft preparation, review and editing)

## Acknowledgements

The authors are grateful to clinical staff in CAMEO, Cambridgeshire and Peterborough NHS Trust, for help with participant recruitment.

## Appendix Expectation-Maximization

Expectation-Maximization adjusts the participant-level parameter estimates and the group-level parameter estimates to make them maximally consistent with the data (E) and with each other (M). The group level distributions are gamma-shaped. For the purposes of algebraic manipulation, it is convenient to parameterise these in terms of a so-called shape parameter *Κ* and a scale parameter *θ*, but can also they can be equivalently described by their mean *m* = *Κ θ* and variance *σ*^2^ = *Κ θ*^2^. We report values for mean and variance as we believe they are more intuitively understandable by clinical researchers but use the shape-scale notation in this appendix for algebraic clarity. The prior probability for each participant’s parameters under the group parameters in our generative statistical model *G* (not to be confused with *G* for ‘gold’) is:

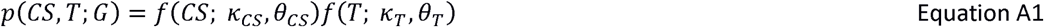

At the participant level, we denote the (empirical) posterior beliefs about the parameters of each participant *j,* furnishing data *d*_*j*_ as Q[T, *CS*; *d*_*j*_]. Again *Q* here not to be confused with action values. We write *v* = {*T,CS*} and the probability that the participant made all the decisions they did under equation 5 as P[*d*_*j*_ | *v*; *G*]. Now, we seek to maximise the aforementioned consistency by minimizing the free energy of the entire model, given the data:

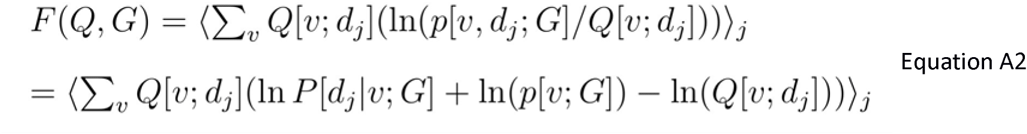

Substituting Eq. 1a gives

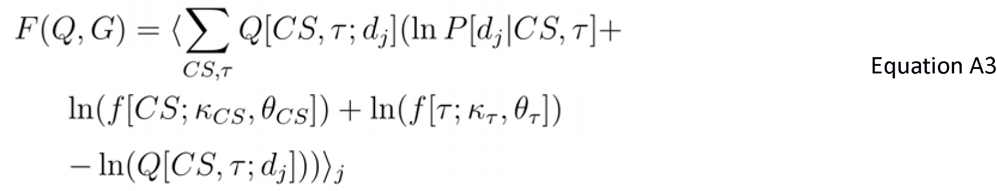

In the maximisation phase, we keep the form of all Q[T, *CS*; *d*_*j*_] fixed and differentiate Eq. A3 w.r.t. *G*, that is, *κ*_*CS*_,*θ*_*CS*_,*κ*_*T*_,*θ*_*T*_. We can derive:

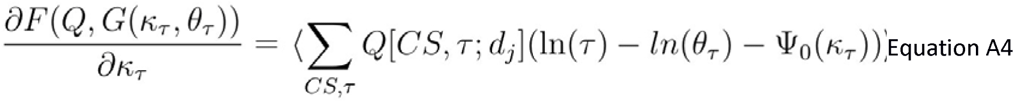

Where Ψ_0_ is the digamma function. In an analogous fashion,

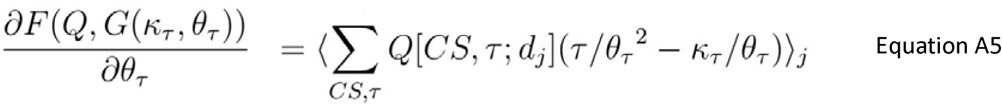

Setting Eq. A5 equal to zero and using integrals instead of sums, as the parameters are continuous, gives for the group mean *T*_*m*_:

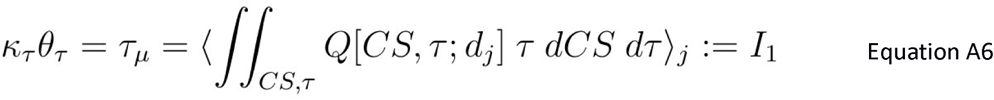

We evaluated this expectation numerically and used it to express *κ*_*T*_ = *I*_1_/ *θ*_*T*_. We then set Eq. A4 equal to zero and obtained:

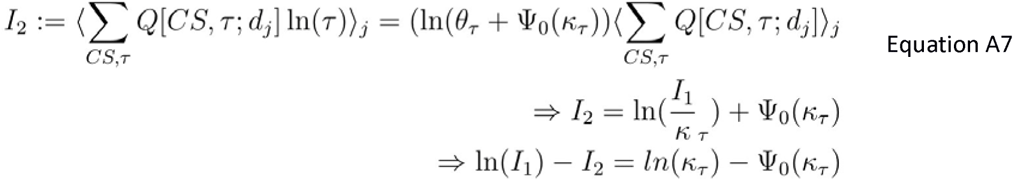

Again, *I*_2_ is integrated numerically and Eq. A7 solved for *κ*_*T*_ by refining its standard approximate algebraic solution numerically.

The Expectation step can proceed much more straightforwardly, in terms of algebra, as Bayes theorem gives:

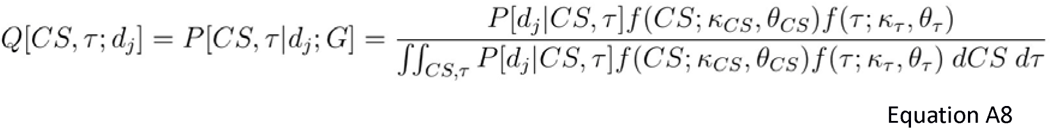

In this low-dimensional parameter space the denominator was numerically tractable using Monte-Carlo integration, but for high dimensional models algebraic approximations would be needed.

Once EM has converged, equation A8 gives the probability distributions for individual participants, which can themselves be expressed in terms of shape and scale, or mean and variance.

